# Leveraging Hardy–Weinberg disequilibrium for association testing in case-control studies

**DOI:** 10.1101/2020.11.14.382796

**Authors:** Lin Zhang, Lei Sun

## Abstract

In a case-control association study, deviation from Hardy-Weinberg equilibrium (HWE) or Hardy-Weinberg dis-equilibrium (HWD) in the control group is usually considered as evidence for potential genotyping error, and the corresponding SNP is then removed from the study. On the other hand, assuming HWE holds in the study population, a truly associated SNP is expected to be out of HWE in the case group. Efforts have been made in combining association tests with tests of HWE in the cases to increase the power of detecting disease susceptibility loci (Song and Elston (2006), Wang and Shete (2010)). However, these existing methods are ad-hoc and sensitive to model assumptions. Utilizing the recent robust allele-based (RA) regression model for conducting allelic association tests (Zhang and Sun (2020)), here we propose a joint RA test that naturally integrates association evidence from the traditional association test and a test that evaluates the difference in HWD between the case and control groups. The proposed test is robust to genotyping error, as well as to potential HWD in the population attributed to factors that are unrelated to phenotype-genotype association. We provide the asymptotic distribution of the proposed test statistic so that it is easy to implement, and we demonstrate the accuracy and efficiency of the test through extensive simulation studies and an application.

## 1 Introduction

In a standard case-control study, deviation from Hardy–Weinberg equilibrium (HWE) or Hardy–Weinberg dis-equilibrium (HWD) in the control group is usually considered as evidence for potential genotyping error, and the corresponding SNP is then removed from the study. On the other hand, assuming HWE holds in the study population, a truly associated SNP is expected to be out of HWE in the case group.

Several authors have proposed methods that combine the traditional association test with a test of HWE in the case sample to increase the power of detecting disease susceptibility loci. For example, Song and Elston (2006) proposed the ‘weighted average (WA) statistic’, which is a weighted sum of the Cochran–Armitage (CA) trend association test statistic and the HWD trend test statistic, and the weights are determined by the magnitudes of the observed test statistics. Wang and Shete (2008) proposed two approaches that aggregate the p-values of likelihood ratio test for association and an exact test of HWE in the case sample using the tail-strength measure (Taylor and Tibshirani, 2006). However, these existing methods are ad-hoc and sensitive to genetic model assumptions.

We have earlier proposed a new allele-based (RA) regression model, that i) analyzes alleles instead of genotypes (i.e. paired alleles), and ii) treats alleles as the response variable, which allows for explicit modeling of HWD. We showed that, for a binary phenotype, the score test statistic derived from the proposed RA method encompasses the classical allelic association test as a special case.

Utilizing the RA regression framework, here we propose a RA joint test that naturally integrates the traditional association evidence with evidence from the *difference* in HWD between the case and control samples. The proposed test is robust to genotyping error, as well as to potential population level HWD attributed to factors that are unrelated to phenotype-genotype association. We provide the asymptotic distribution of the proposed test statistic under the null hypothesis of no association, and we demonstrate the accuracy and efficiency of the test through extensive simulation and application studies.

## 2 Revisit the Robust Allele-based (RA) association test

The RA framework partitions a genotype as

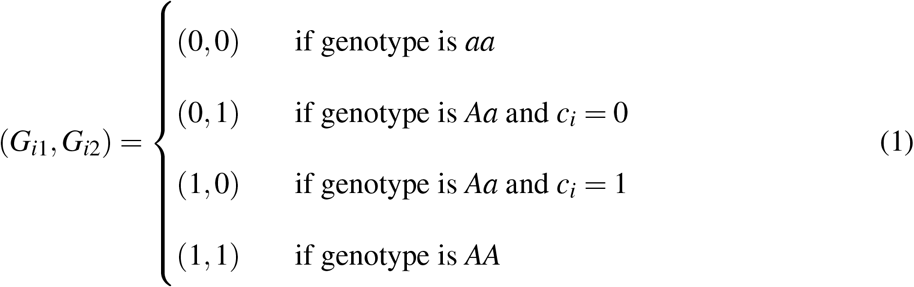

where 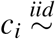 Bernoulli(1/2). The term *c*_*i*_ ensures the distributional assumption of the alleles; see Section **??** for details.

Assume that we have an independent sample of size *n*. For a given biallelic SNP *G* and a phenotype trait *Y* of interest, the RA regression framework for association analysis is

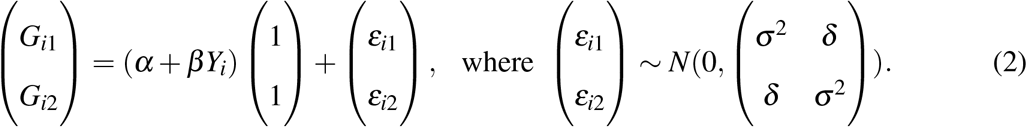

Based on (2), testing

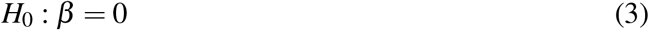

evaluates the association between the SNP and phenotype of interest.

The RA regression (2) provides a unifying framework for allele-based association testing. In the case of a binary outcome, the score test statistic of *H*_0_ : *β* = 0 is

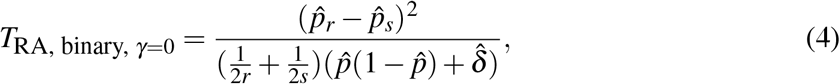

where 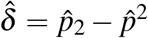 is a sample estimate of the measure of HWD under *H*_0_. If we assume HWE holds in the population (i.e. *δ* = 0), *T*_RA, binary, *γ*=0, *δ* =0_ is identical to the classical allelic test *T*_allelic_ in (**??**).

So far, the measure of HWD, *δ* in (2), is treated as a nuisance parameter and estimated under the null hypothesis of no association. However, for a truly associated SNP, *δ* is expected to be different between the case and control samples, which we illustrate in the next section.

## 3 Hardy–Weinberg disequilibrium in case and control samples

Prior to conducting a case-control genome-wide association study (GWAS), testing HWE in the control sample and excluding SNPs out of HWE is a routine part of data quality control. However, there is no universal agreement on the HWE screening p-value threshold, and it is also not clear if HWE should be evaluated in the case sample as well. For example, Anderson et al. (2010) argues that HWE should be evaluated in the control sample only and the p-value threshold can range from 0.001 to 10^−7^ in many GWAS. On the other hand, Marees et al. (2018) suggests screening out SNPs with test of HWE p-values less than 10^−6^ in controls and 10^−10^ in cases. For largescale population-based GWAS, the p-value threshold is much smaller. The UK Biobank (with total sample size of ~500,000) genotyping quality control procedure adopts a HWE p-value threshold of 10^−12^ per ~4,000 sample batch (Bycroft et al., 2018).

The (implicit) assumption of the HWE-based screening practice is that, without genotype error, HWE holds in the control population or HWD is not detectable in a control sample. However, for a truly associated SNP, HWD may be expected in either the case or control populations due to true association. Recall the example that examines the association between HLA-DQ3 with cervical intra-epithelial neoplasia (Sasieni, 1997). The test of HWE yields a p-value of 3.54 × 10^−10^ in the controls and 0.0387 in the cases; see Table 1 for details. The HLA-DQ3 marker would have been removed from the study following the standard HWE-based screening practice, but HLA-DQ3 is well-known to be associated with cervical intra-epithelial neoplasia (Apple et al., 1994).

**Table 1:** The HLA-DQ3 example that studies women with cervical intra-epithelial neoplasia 3.

|  | Genotype Counts |  |  |  | The HLA-DQ3 example |  |  |  |  |  |  |
| --- | --- | --- | --- | --- | --- | --- | --- | --- | --- | --- | --- |
| | *aa* | *Aa* | *AA* | Total | *aa* | *Aa* | *AA* | Total | $\hat{p}$ | $\hat{\delta}$ | $p_{\text{HWE}}$ |
| Case | $r_0$ | $r_1$ | $r_2$ | $r$ | 40 | 45 | 28 | 113 | 0.4469 | 0.0481 | 0.0387 |
| Control | $s_0$ | $s_1$ | $s_2$ | $s$ | 273 | 100 | 43 | 416 | 0.2236 | 0.0534 | $3.54 \times 10^{-10}$ |
| Total | $n_0$ | $n_1$ | $n_2$ | $n$ | 313 | 145 | 71 | 529 | 0.2713 | 0.0606 | $1.74 \times 10^{-12}$ |

Before providing simulated examples, we first offer some theoretical insights on the departure from HWE in the case or control populations. Suppose a truly associated SNP *G* is in HWE at the population level, with risk allele frequency *p*; see Table **??** for definitions. The genotype frequencies of (*aa, Aa, AA*) are then {(1 − *p*)^2^, 2(1 − *p*)*p, p*^2^}. In this work, we still use *A* to denote the risk allele, but the risk allele can be either the minor allele with frequency ≤ 0.5 or the major allele with frequency > 0.5, and thus *p* can range from 0 to 1. Recall the definition of disease penetrance probabilities,

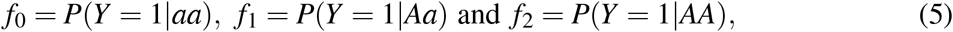

and disease prevalence,

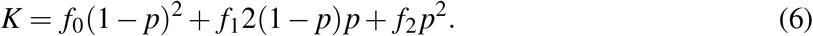

Thus, the genotype frequencies of *AA* and *Aa* in the case population are

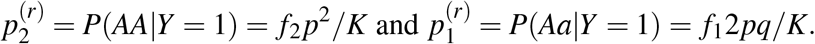

The frequency of allele *A* in the case population is

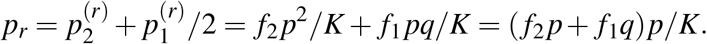

Thus, we expect departure from HWE in the case population, and the amount HWD is

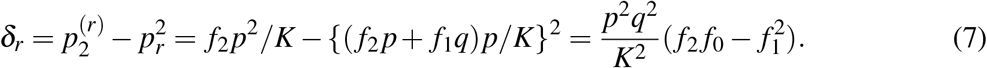

Similarly, the amount of HWD in the control population is

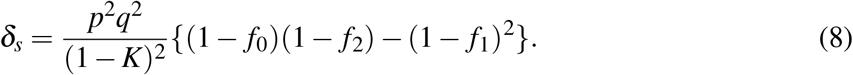

Results in (7) and (8) show that, for a truly associated SNP, even if HWE holds at the population level and there is no genotype error, HWE may not hold in the case or control populations. The magnitude and the direction of HWD in case and control samples are determined by the underlying genetic model, which specifies the relationship between *f*_0_, *f*_1_ and *f*_2_. The sampling scheme matters as well, but for the purpose of this study, we assume random sampling of a case and a control, respectively, from the case and control populations.

We now illustrate the expected HWD, in the absence of genotyping error, in case and control samples with two simulated examples. Suppose a truly associated SNP, *G*, is in HWE at the population level with risk allele frequency *p* = 0.2. We randomly sample genotypes (*aa, Aa, AA*) based on probabilities {0.64, 0.32, 0.04}, and assign them to the case group with probability *f*_*k*_ and the control group with probability (1 − *f*_*k*_), where *k* = 0, 1 and 2 refer to the number of risk allele *A*. The sampling process stops when we have *r* =1,000 cases and *s* =1,000 controls. Table 2 summarizes a set of simulated genotype counts and the corresponding sample estimates under an additive genetic model (*f*_1_ = ( *f*_0_ + *f*_2_)/2) and a recessive genetic model ( *f*_0_ = *f*_1_ < *f*_2_).

**Table 2:** Two illustrative examples of HWD in case and control samples for a truly associated SNP, assuming the SNP is in HWE at the population level without genotyping error. The risk allele frequency *p* = 0.2. Top panel: additive genetic model assuming *f*_1_ = (*f*_0_ + *f*_2_)/2 with *f*_0_ = 0.08, *f*_1_ = 0.10 and *f*_2_ = 0.12. Bottom panel: recessive genetic model assuming *f*_0_ = *f*_1_ < *f*_2_ with *f*_0_ = *f*_1_ = 0.08 and *f*_2_ = 0.12.

| Additive genetic model |  |  |  |  |  |  |  |  |  |
| --- | --- | --- | --- | --- | --- | --- | --- | --- | --- |
| | *aa* | *Aa* | *AA* | Total | $\hat{p}$ | $\hat{\delta}^{\text{ADD}}$ | $p_{\text{HWE}}$ | $p$ | $\delta^{\text{ADD}}$ |
| Case | 566 | 390 | 44 | 1000 | 0.2390 | **-0.0131** | **0.0225** | 0.2364 | $-1.32 \times 10^{-3}$ |
| Control | 663 | 306 | 31 | 1000 | 0.1840 | **-0.0029** | **0.5475** | 0.1965 | $-1.23 \times 10^{-5}$ |
| Total | 1229 | 696 | 75 | 2000 | 0.2115 | **-0.0072** | **0.0524** | 0.2000 | 0 |

Table 2: **Two illustrative examples of HWD in case and control samples for a truly associated SNP, assuming the SNP is in HWE at the population level without genotyping error.**
| Recessive genetic model |  |  |  |  |  |  |  |  |  |
| --- | --- | --- | --- | --- | --- | --- | --- | --- | --- |
| | *aa* | *Aa* | *AA* | Total | $\hat{p}$ | $\hat{\delta}^{\text{REC}}$ | $p_{\text{HWE}}$ | $p$ | $\delta^{\text{REC}}$ |
| Cases | 632 | 306 | 62 | 1000 | 0.2150 | **0.0158** | **0.0031** | 0.2157 | 0.0123 |
| Control | 606 | 349 | 45 | 1000 | 0.2195 | **-0.0032** | **0.5572** | 0.1986 | -0.0011 |
| Total | 1238 | 655 | 107 | 2000 | 0.2173 | **0.0063** | **0.0974** | 0.2000 | 0 |

Under the additive genetic model where *f*_1_ = (*f*_0_ + *f*_2_)/2, (7) and (8) can be further simplified to

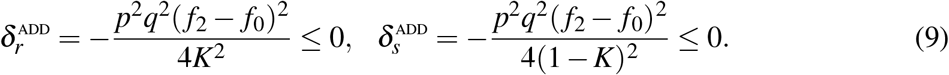

Thus, if the SNP is truly associated, i.e. *f*_2_ ≠ *f*_0_, 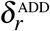 and 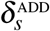 are both negative. The negative sample estimates of *δ*_*r*_ and *δ*_*s*_ in Table 2 verify the theoretical derivations in (9). The test of HWE is rejected in the case sample but not in the control sample at *α* = 0.05.

If the disease prevalence *K* is small, based on (9), the HWD in the control population, attributed to true phenotype-genotype association, is too small to be detected using a control sample of moderate sample size at a stringent *α* level. Hence, if the genetic model is additive and the SNP is in HWE at the population level, the HWD in the control population is expected to be negligible without genotyping error. This may explain why testing for HWD in the control sample has been used for detecting for potential genotyping error, as part of data quality control.

The second simulated example assumes a recessive genetic model where *f*_0_ = *f*_1_ < *f*_2_, with all the other settings remain the same as the first example. Under a recessive genetic model, (7) and (8) can be further simplified to

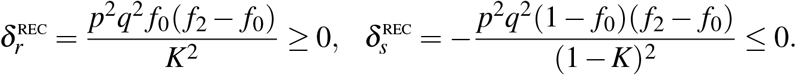

For a truly associated SNP *G*, 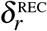 and 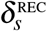 are in opposite directions, which is verified by the sample estimates 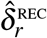 and 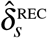 in Table 2.

Neither of the two simulated examples would exclude the truly associated SNP from analysis, given the sample size and genetic effect size. However, in practice, the HWE-based screening practice is becoming more problematic as the sample size of GWAS continues to grow. For example, the sample size of the UK Biobank data is ~ 500,000 (Bycroft et al., 2018). In that case, if *s*=250,000 and 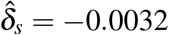 as in the simulated recessive example, the p-value of testing HWE in the control sample would be 8.92 × 10^−24^. This truly associated SNP may be considered as out of HWE due to genotying error, and falsely removed from the study. Further to this consideration, the two examples here also suggest that, rather than treating the HWD measure *δ* as a nuisance parameter, a better method should utilize the expected HWD in the control sample (and the case sample) attributed to true association.

We thus propose a unifying and flexible association test, RA joint test, that can be more powerful than the traditional association testing, by incorporating evidence for difference in HWD between the case and control samples. Testing *δ*_*r*_ = *δ*_*s*_ instead of *δ*_*s*_ = 0 (i.e. HWE holds in the control sample) also provides robustness to departure from HWE in a sample due to genotyping error or at the population level due to factors other than phenotype-genotype association.

## 4 A robust allele-based (RA) joint test for case-control studies

To develop the RA joint association association test for case-control studies, we adopt the general RA regression framework in (2) but handle the *δ* parameter differently. For a case-control study with independent samples, the RA joint association test is derived from

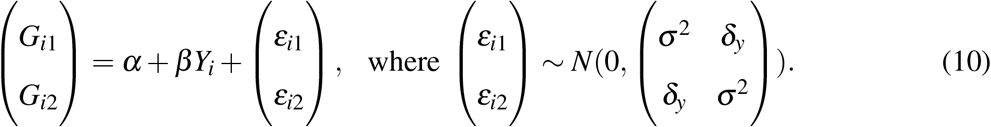

Unlike the original RA framework in (2) that assumes *δ*_*y*_ being identical for all samples, the RA joint test allows *δ*_*y*_ to be different for the case and control groups,

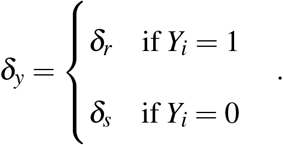

The RA joint association test evaluates

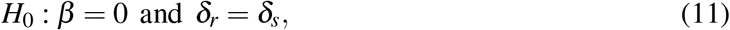

and estimates *δ*_*r*_ and *δ*_*s*_ under the alternative hypothesis. We emphasize that in (11) we test if HWD differs between the case and control samples, rather than if HWE holds in either the case or control samples, to gain robustness to HWD due to genotype error or HWD at the population level.

The RA joint test statistic is

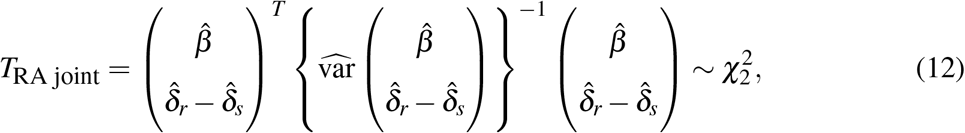

where

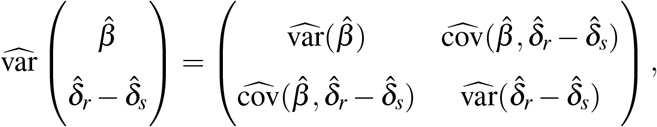

and 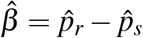.

We decompose *T*_RA joint_ into three components to provide additional analytical insights: (i) *T*_*p*_ captures the differences in risk allele frequencies between the case and control samples; (ii) *T*_*δ*_ captures the difference in HWD estimates between the case and control samples; (iii) *T*_*p,δ*_ adjusts for the correlation between *T*_*p*_ and *T*_*δ*_,

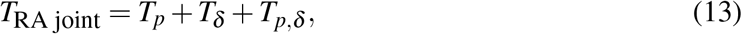

where

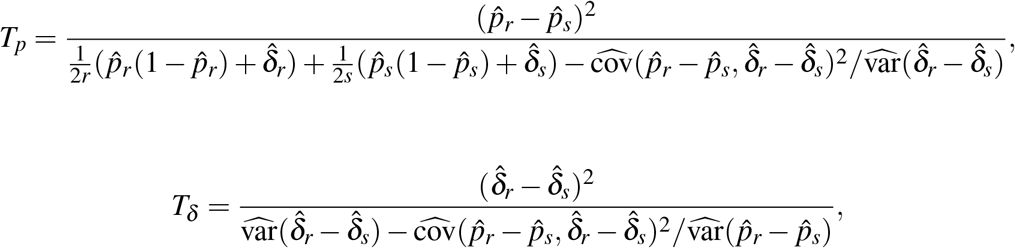

and

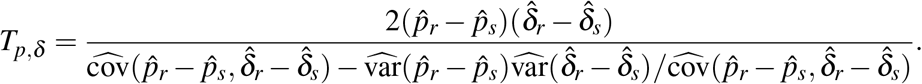

Note that *T*_*p*_ is similar to *T*_allelic, Schaid_ in (**??**) (Schaid and Jacobsen, 1999), except that *T*_*p*_ subtracts 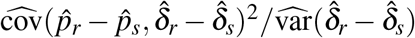 in the denominator to account for the dependence between *T*_*p*_ and *T*_*δ*_.

Assume the case and control samples are independent of each other as in a standard case-control study, it is easy to show that 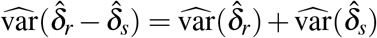, 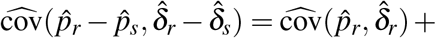 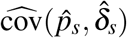, and 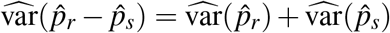, where the variance-covariance components can be estimated in the case and control samples separately.

For the case sample, as the sample size *r* → ∞, the asymptotic distribution of 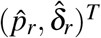 is

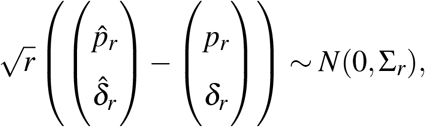

where

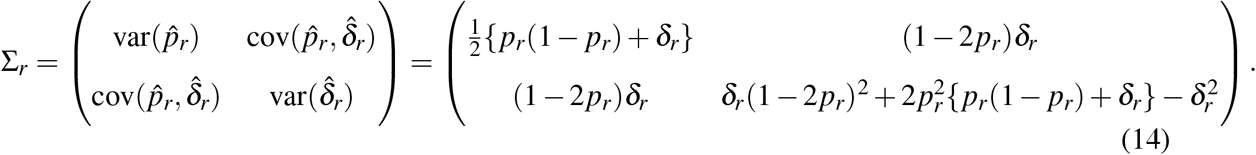

Σ_*s*_ is defined similarly by replacing *r* in (14) with *s*; see Supplementary Material **??** for the detailed derivation.

The variance-covariance matrix of 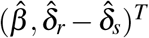 can then be written as

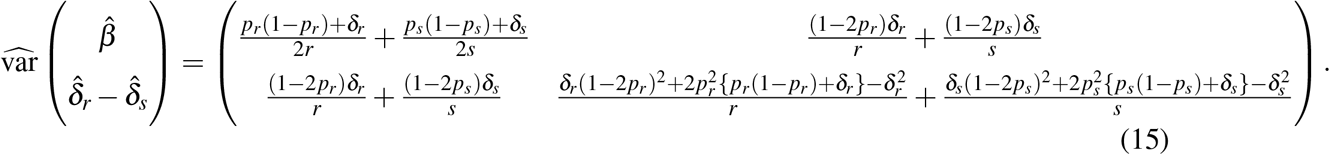

Another major distinction between the original RA test proposed and the RA joint test studied here is that the former uses a score test and estimates all the parameters under the null hypothesis of no association, while the latter uses the Wald’s test and estimates the parameters under the alternative hypothesis. This is because by estimating the parameters under the alternative, the proposed RA joint test is able to leverage the difference between 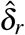 and 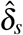 to gain additional power for association testing.

Before diving into the performance evaluation of the proposed RA joint test, we first review another commonly used 2 d.f. genetic association test, the traditional genotypic test, which expands our method comparison from the traditional 1 d.f. association test to 2 d.f.; see Table **??** for background on the different types of genetic association tests.

## 5 The traditional prospective association tests

In the traditional phenotype-on-genotype or prospective association testing framework, two commonly used tests are the 1 d.f. additive test and the 2 d.f. genotypic test.

The traditional 1 d.f. additive test codes the genotypes *aa*, *Aa* and *AA* as 0, 1 and 2, respectively.

The additive test for a case-control study uses logistic regression,

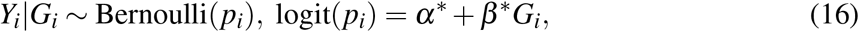

and evaluates the association between *Y* and *G* by testing

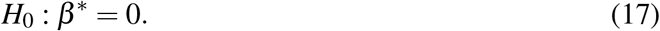

The traditional 2 d.f. genotypic test treats the three genotypes *aa*, *Aa* and *AA* as categories. The genotypic test for a case-control study also uses logistic regression,

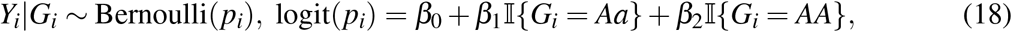

where, without loss of generality, genotype *aa* is chosen to be the baseline category,

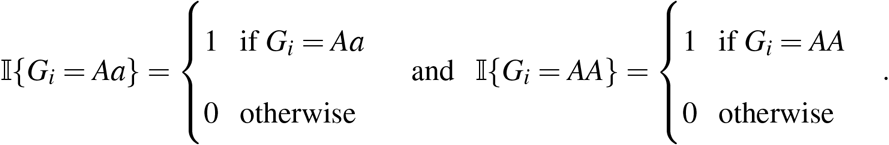

Testing

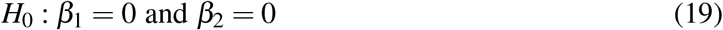

evaluates the association between *Y* and *G*.

For fair comparison with the proposed RA joint test, we evaluate both (17) and (19) using the Wald’s test, denoted as *T*_additive_ and *T*_Genotypic_, respectively. Conceptually, because both the additive and genotypic tests analyze genotypes directly (i.e. not ‘decoupling’ the two paired alleles), they are robust to departure from HWE. However, the corresponding HWD adjustments are not well understood analytically. We refer readers to Supplementary Materials for additional discussion of these two tests, and here we move on to method comparison.

## 6 Simulation studies

In this section, we use simulated data to evaluate the proposed RA joint test and compare it with the traditional 1 d.f. additive test and the 2 d.f. genotypic test. We first assess the empirical type 1 error rate and the power of the three tests at *α* = 0.05 as in a candidate gene study, and we fix the case and control sample size at *r* = *s* = 500. We further investigate the difference between the two 2 d.f. tests at the GWAS nominal rate of *α* = 5 × 10^−8^ (Dudbridge and Gusnanto, 2008), with a larger sample size *r* = *s* =2,500 as achieved by many existing GWAS.

### 6.1 Simulation setup

Suppose the risk allele frequency of *G* is *p* with population level departure from HWE, *δ* = *p*_2_ − *p*^2^.

The population level genotype frequencies of *aa*, *Aa* and *AA* are

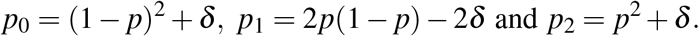

We follow the same sampling procedure used for the two illustrative examples in Section 3, but without the assumption of HWE at the population level. We sample the genotypes based on probabilities {*p*_0_, *p*_1_, *p*_2_}, and each genotype *G* = *k* is assigned to the case group with probability *f*_*k*_ and to the control group with probability (1 − *f*_*k*_), where *f*_*k*_ = *P*(*Y* = 1|*G* = *k*) for *k* = 0, 1 and 2. The sampling procedure stops when there are *r* cases and *s* controls.

### 6.2 Type I error control

We first assess the empirical type I error rate of the proposed RA joint test under the null hypothesis of no association assuming *f*_0_ = *f*_1_ = *f*_2_ = *K* = 0.1. We simulate *r* = *s* = 500 independent samples for both the case and control groups, varying the population risk allele frequency *p* from 0.1 to 0.9 on a grid of 0.1. For each *p*, the departure from HWE, *δ*, takes the value from {0, 0.02, 0.04, 0.06}; when *δ* = 0, HWE holds in the population.

Table 3 summarizes the empirical type I error rate of the proposed RA joint test for each combination of *p* and *δ*, which shows that the proposed test is accurate. Figure 1 shows the histograms and QQ-plots of the 10, 000 p-values when *p* = 0.2 and *δ* = 0, 0.02 and 0.04, providing additional evidence for the accuracy of the proposed test and its robustness to population level HWD.

**Table 3:** Empirical type 1 error rate of the proposed RA joint test at *α* = 0.05 evaluated based on 10,000 simulation replicates for each parameter setting. Each replicate contains a sample of 500 cases and 500 controls.

| Population $\delta$ | Risk allele frequency $p$ | | | | | | | | |
| --- | --- | --- | --- | --- | --- | --- | --- | --- | --- |
|  | 0.1 | 0.2 | 0.3 | 0.4 | 0.5 | 0.6 | 0.7 | 0.8 | 0.9 |
| 0(HWE) | 0.0503 | 0.0496 | 0.0489 | 0.0524 | 0.0485 | 0.0509 | 0.0504 | 0.0495 | 0.0471 |
| 0.02 | 0.0502 | 0.0519 | 0.0548 | 0.0483 | 0.0489 | 0.0513 | 0.0496 | 0.0513 | 0.0524 |
| 0.04 | 0.0502 | 0.0507 | 0.0522 | 0.0501 | 0.0510 | 0.0491 | 0.0493 | 0.0473 | 0.0520 |
| 0.06 | 0.0518 | 0.0503 | 0.0518 | 0.0543 | 0.0507 | 0.0494 | 0.0501 | 0.0484 | 0.0493 |

**Figure 1:**
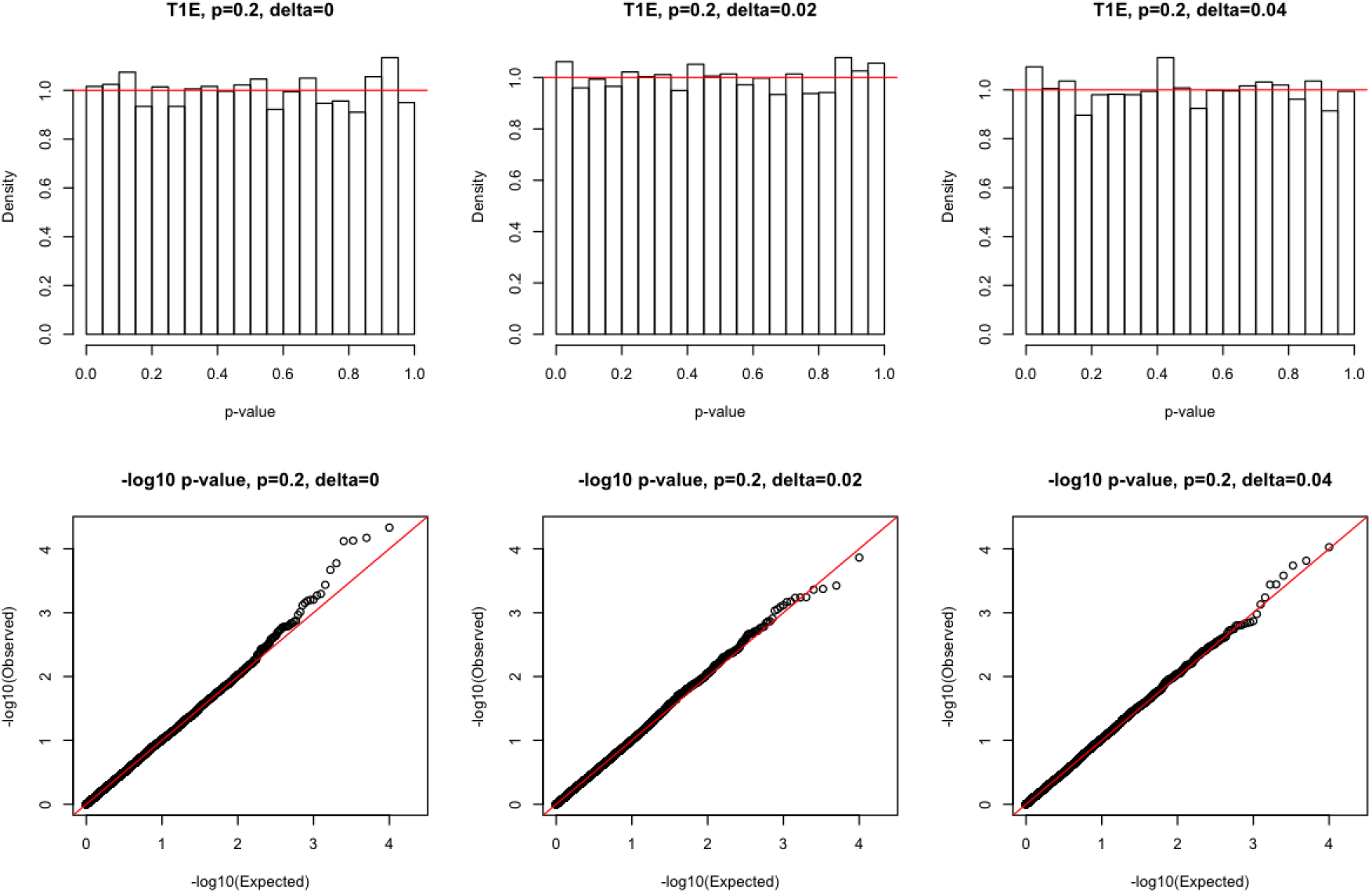
Histograms and QQ-plots of the log_10_ p-values of the proposed RA joint test at *α* = 0.05. The p-values are evaluated based on 10,000 simulation replicates under three settings at *p* = 0.2, with *δ* = 0, 0.02 and 0.04, respectively. Each replicate contains a sample of 500 cases and 500 controls.

### 6.3 Power analysis

We then evaluate the power of the proposed RA joint test, and compare it with the traditional prospective 1 d.f. additive test and the 2 d.f. genotypic test. To examine the impact of genetic models on the power of all three tests, we simulate data following six most common genetic models as summarized in Table 4, each assuming a different relationship between *f*_0_, *f*_1_ and *f*_2_. The risk allele frequency *p* ranges from 0.05 to 0.95, and we here assume HWE holds at the population level. The power is evaluated at *α* = 0.05 based on 1,000 simulation replicates for each parameter setting. Each replicate contains a sample of 500 cases and 500 controls.

**Table 4:** Six common genetic models and their corresponding penetrance specifications for the power studies. For additive, additive on log-scale, recessive, and dominant genetic models, we assume the homozygote relative risk *f*_2_/ *f*_0_ = 1.5 and the corresponding *f*_1_ is determined by the genetic model assumptions; for heterozygous advantage genetic model, we assume the heterozygote relative risk *f*_1_/ *f*_0_ = 1/1.5, and *f*_0_ = *f*_2_; for heterozygous disadvantage genetic model, we assume the heterozygote relative risk *f*_1_/ *f*_0_ = 1.5, and *f*_0_ = *f*_2_.

| Genetic model | Penetrance specifications | | | | heterozygote<br>relative risk<br>($f_1/f_0$) | homozygote<br>relative risk<br>($f_2/f_0$) |
| --- | --- | --- | --- | --- | --- | --- |
| | | $f_0$ | $f_1$ | $f_2$ | | |
| Additive | $f_1 = (f_0 + f_2)/2$ | 0.100 | 0.125 | 0.150 | 1.25 | 1.50 |
| Additive on log-scale | $f_1 = \sqrt{f_0 f_2}$ | 0.100 | 0.122 | 0.150 | 1.22 | 1.50 |
| Recessive | $f_1 = f_0$ | 0.100 | 0.100 | 0.150 | 1.00 | 1.50 |
| Dominant | $f_1 = f_2$ | 0.100 | 0.150 | 0.150 | 1.50 | 1.50 |
| Heterozygous advantage | $f_1 < f_0 = f_2$ | 0.150 | 0.100 | 0.150 | 0.67 | 1.00 |
| Heterozygous disadvantage | $f_1 > f_0 = f_2$ | 0.150 | 0.225 | 0.150 | 1.50 | 1.00 |

Under the additive genetic model, as expected, the proposed RA joint test and the traditional genotypic test have lower power than the traditional additive test (Figure 2a). However, this loss can be capped at 11% based on a theoretical comparison between 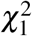 and 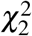 distributions, regardless of the genetic effect size, sample size and the nominal *α* level (Chen et al., 2018). The power under the additive on log-scale genetic model, or the multiplicative genetic model, shows a similar pattern (Figure 2b). This is because when *f*_2_ and *f*_2_/ *f*_0_ are fixed, 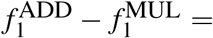 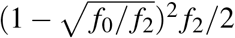 and the difference is usually close to 0. For example, when *f*_2_ = 0.15 and *f*_2_/ *f*_0_ = 1.5 for both additive and multiplicative genetic models, 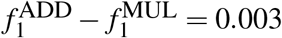.

**Figure 2:**
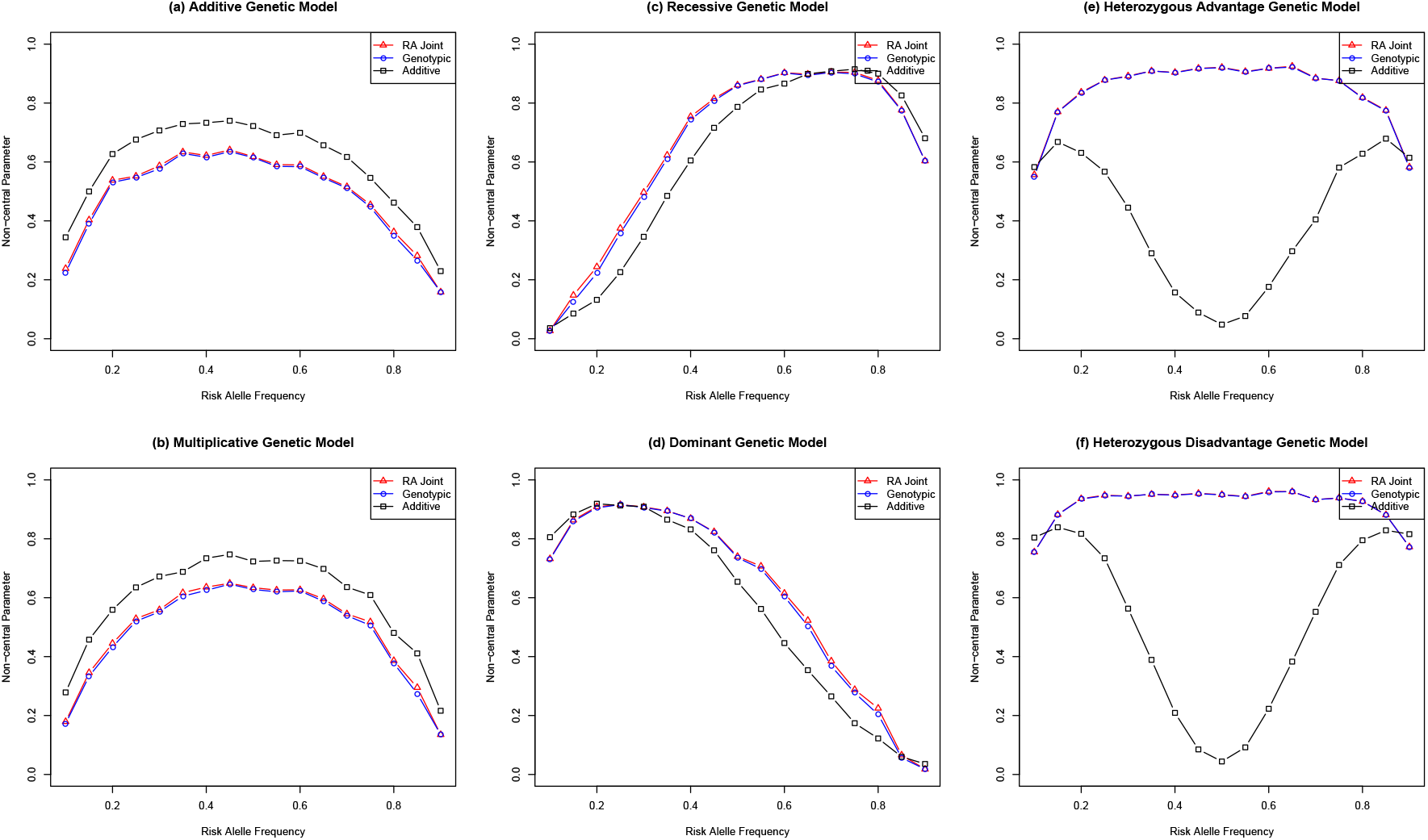
Power of *T*_RA joint_, *T*_genotypic_ and *T*_additive_ under six common genetic models. The penetrance probabilities for all the genetic models are summarized in Table 4. The empirical power is evaluated with 1, 000 replicates at *α* = 0.05. We sample *r* = *s* = 500 for both case and control groups, and assume HWE holds at the population level, i.e. *δ* = 0. Red triangles 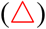 symbolize the proposed RA joint test; blue circles 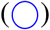 symbolize the traditional 2 d.f. genotypic test; black squares (◻) symbolize the traditional 1 d.f. additive test.

On the other hand, under the recessive and dominant genetic models, the proposed RA joint test and the traditional genotypic test have slightly higher power than the traditional additive test (Figures 2c and 2d), and under the heterozygous advantage and disadvantage models, the two 2 d.f. tests have substantially higher power than the 1 d.f. additive test (Figures 2e and 2f). For example, for the heterozygous disadvantage model when *f*_0_ = *f*_2_ = 0.15 and *f*_1_ = 0.225 and risk allele frequency *p* = 0.5, the power of the two 2 d.f. tests are close to 100% while the power of the 1 d.f. additive test is close to 5%. This is because at *p* = 0.5, the genotype frequencies are (0.3125, 0.3750, 0.3125) for the case population and (0.2431, 0.5139, 0.2431) for the control population, which leads to *p*_*r*_ = *p*_*s*_ = 0.5. Although the additive test does not directly test allele frequency being the same, its test statistic essentially only captures the difference between the case and control allele frequencies using an adjusted variance.

The power study in Figure 2 assumes HWE holds in the population. We show in Supplementary Figure **??** that, as the population level HWD (*δ*) increases, the power of the traditional 1 d.f. additive test drops while that of the traditional genotypic test and the proposed RA joint test increases. That is, under the (linear and multiplicative) additive models, the power loss of the RA joint test diminishes as *δ* gets larger, while under the other genetic models, the power gain of the RA joint test increases as *δ* gets larger. We further verify the empirical findings using theoretical power computed under the same settings with or without the HWE assumption; see Supplementary Figures **??** and **??**.

### 6.4 Type I error and power using the GWAS nominal rate of *α* = 5 × 10^−8^

So far, we have examined the empirical type 1 error rate and power of the traditional 1 d.f. additive test, 2 d.f. genotypic test and the proposed RA joint test at the nominal rate *α* = 0.05. In modern genome-wide association studies where tens of millions SNPs are tested, the commonly used nominal rate is *α* = 5 × 10^−8^ to adjust for multiple hypothesis testing. We have shown in Section 6.3 that the power difference is clear between the 1 d.f. additive test and the two 2 d.f. tests under the six genetic models, so we restrict our attention here to investigate the difference between *T*_RA joint_ and *T*_genotypic_.

To assess the type 1 error rate of the two d.f. association tests at the GWAS significance level, we follow the same simulation setting as in Section 6.2, except that we use *α* = 5 × 10^−8^ and increase the sample size to *r* = *s* =2,500 for both the case and control groups. The empirical type 1 error is evaluated based on 10^10^ replicates for each parameter setting. Table 5 summarizes the empirical type 1 error rate for each *p* and *δ* combination. Overall, the proposed RA joint test has well-controlled type 1 error rate and is robust to departure from HWE at the population level. When the risk allele frequency is very low or very high (e.g. 0.1 or 0.9), the test is slightly conservative. On the other hand, the traditional 2 d.f. genotypic test appears to be generally conservative at the *α* = 5 × 10^−8^ level, which has not been noted before to the best of our knowledge.

**Table 5:** Empirical type 1 error rates of the proposed RA joint test and the traditional genotypic test at the GWAS nominal rate of. *α* = 5 × 10^−8^, evaluated based on 10^10^ simulation replicates for each parameter setting. Each replicate contains a sample of 2,500 cases and 2,500 controls.

| The proposed 2 d.f. RA joint test |  |  |  |  |  |
| --- | --- | --- | --- | --- | --- |
| Population<br>$\delta$ | Risk allele frequency $p$ | | | | |
|  | 0.1 | 0.3 | 0.5 | 0.7 | 0.9 |
| 0(HWE) | $3.19 \times 10^{-8}$ | $5.50 \times 10^{-8}$ | $5.23 \times 10^{-8}$ | $5.43 \times 10^{-8}$ | $3.29 \times 10^{-8}$ |
| 0.02 | $4.69 \times 10^{-8}$ | $5.13 \times 10^{-8}$ | $5.85 \times 10^{-8}$ | $5.23 \times 10^{-8}$ | $4.74 \times 10^{-8}$ |
| 0.04 | $5.18 \times 10^{-8}$ | $5.78 \times 10^{-8}$ | $5.56 \times 10^{-8}$ | $5.47 \times 10^{-8}$ | $5.02 \times 10^{-8}$ |
| 0.06 | $5.25 \times 10^{-8}$ | $5.77 \times 10^{-8}$ | $5.56 \times 10^{-8}$ | $5.48 \times 10^{-8}$ | $4.80 \times 10^{-8}$ |
| The traditional 2 d.f. genotypic test |  |  |  |  |  |
| Population<br>$\delta$ | Risk allele frequency $p$ | | | | |
|  | 0.1 | 0.3 | 0.5 | 0.7 | 0.9 |
| 0(HWE) | $1.67 \times 10^{-8}$ | $4.26 \times 10^{-8}$ | $4.17 \times 10^{-8}$ | $4.05 \times 10^{-8}$ | $1.90 \times 10^{-8}$ |
| 0.02 | $2.51 \times 10^{-8}$ | $3.88 \times 10^{-8}$ | $4.64 \times 10^{-8}$ | $3.99 \times 10^{-8}$ | $2.62 \times 10^{-8}$ |
| 0.04 | $3.05 \times 10^{-8}$ | $4.42 \times 10^{-8}$ | $4.55 \times 10^{-8}$ | $4.27 \times 10^{-8}$ | $2.66 \times 10^{-8}$ |
| 0.06 | $2.81 \times 10^{-8}$ | $4.62 \times 10^{-8}$ | $4.58 \times 10^{-8}$ | $4.14 \times 10^{-8}$ | $2.49 \times 10^{-8}$ |

We then assess the power of the proposed RA joint test and the traditional 2 d.f. genotypic test. We follow the same simulation setting as in Section 6.3 using the six common genetic models; see Table 4 for details. We fix *f*_2_ = 0.1 and vary the heterozygote relative risk or the homozygote relative risk to reflect different genetic effect sizes. For the heterozygous advantage genetic model, the heterozygote relative risk (*f*_1_/ *f*_0_) takes values from {1/1.6, 1/1.5, 1/1.4}; for the heterozygous disadvantage genetic model, the heterozygote relative risk ( *f*_1_/ *f*_0_) takes values from {1.6, 1.5, 1.4}; for the other genetic models, the homozygote relative risk ( *f*_2_/ *f*_1_) takes values from {1.6, 1.5, 1.4}. The risk allele frequency *p* ranges from 0.05 to 0.95, and we here assume HWE holds at the population level; see Supplementary Figures **??** for the power under HWD. The power is evaluated at *α* = 5 × 10^−8^ based on 1,000 simulation replicates for each parameter setting. Each replicate contains a sample of 2,500 cases and 2,500 controls.

Figure 3 plots the empirical power under the six genetic models with varying degrees of genetic effect size. Although the power of *T*_RA joint_ and *T*_genotypic_ are not identical, they generally follow the same pattern as seen in Section 6.3 when the nominal rate was *α* = 0.05 and sample size was 500 cases and 500 controls.

**Figure 3:**
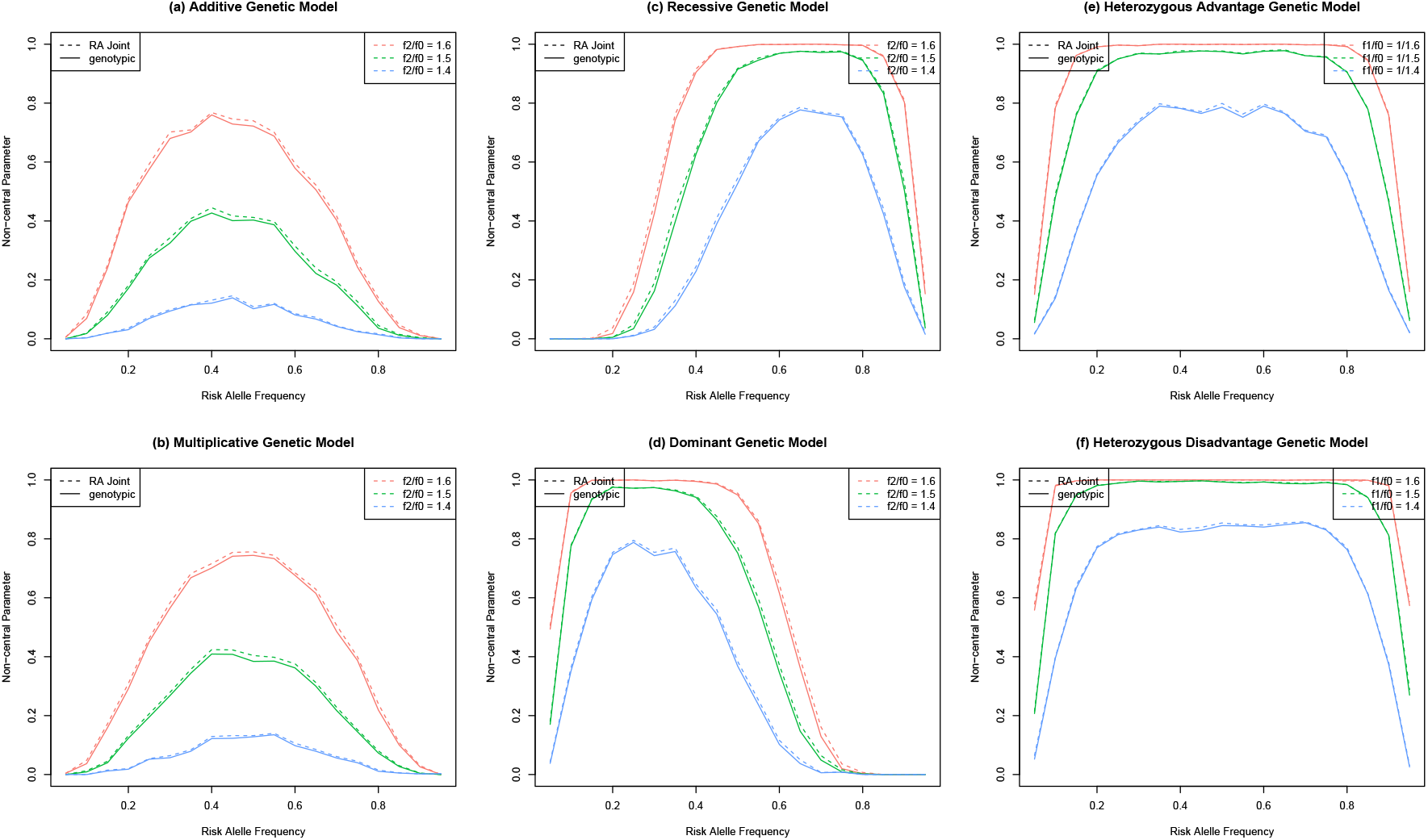
Empirical power of *T*_RA joint_ and *T*_genotypic_ under six common genetic models. *f*_2_ is fixed at 0.1. For the heterozygous advantage genetic model, the heterozygote relative risk (*f*_1_/ *f*_0_) takes values from {1/1.6, 1/1.5, 1/1.4}; for the heterozygous disadvantage genetic model, the heterozygote relative risk ( *f*_1_/ *f*_0_) takes values from {1.6, 1.5, 1.4}; for the other genetic models, the homozygote relative risk ( *f*_2_/ *f*_1_) takes values from {1.6, 1.5, 1.4}. The empirical power is evaluated with 1, 000 replicates at *α* = 5 × 10^−8^. We sample *r* = *s* =2,500 for both case and control groups, and assume HWE holds at the population level, i.e. *δ* = 0. The dashed curves represent the proposed RA joint test; the solid curves represent the traditional 2 d.f. genotypic test.

In this section, we have demonstrated that the proposed RA joint test is accurate for *α* = 0.05 and *α* = 5 × 10^−8^. We also examine its power under six common genetic models, and compare it with the traditional 1 d.f. additive test and the 2 d.f. genotypic test. The power difference between the proposed RA joint test and the traditional 1 d.f. additive test is evident under the six genetic models. However, it appears that the power of *T*_RA joint_ is practically identical to that of *T*_genotypic_. In the next section, we provide theoretical insights on the power of the two 2 d.f. association tests by comparing their non-central parameters (NCPs), and conclude that *T*_RA joint_ and *T*_genotypic_ are indeed different tests.

## 7 Theoretical power comparison of *T*_RA joint_ and *T*_genotypic_ using the GWAS significance level of *α* = 5 × 10^−8^

In this section, we examine the theoretical power of the two tests through non-central parameters. We first show that the general form of the non-central parameter of *T*_genotypic_ is

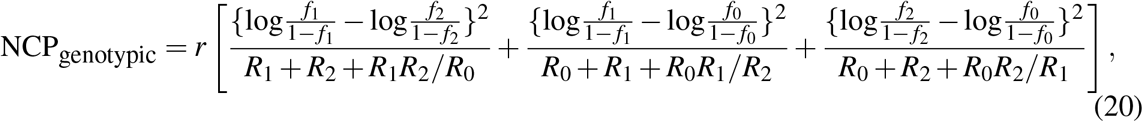

where

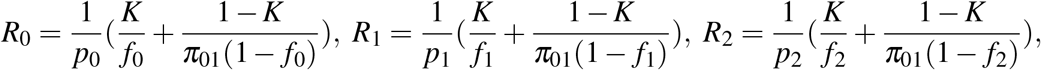

and *π*_01_ = *s/r* is the ratio of the number of controls to the number of cases. Due to the complex form of NCP_RA joint_, we direct the readers to Supplementary Material **??** for a full derivation of NCP_RA joint_.

Figure 4 plots NCP_RA joint_ and NCP_genotypic_ against the risk allele frequency under six common genetic models. Under both the additive and multiplicative genetic models (Figures 4a and 4b), NCP_RA joint_ can be substantially larger than NCP_genotypic_ as the effect size increases (i.e. from blue, green to red). For the dominant (Figure 4c) and recessive (Figure 4d) genetic models, when the effect size is large, e.g. *f*_2_/ *f*_0_ = 2.25 (red curves), NCP_RA joint_ can exceed NCP_genotypic_ by a consequential margin. For example, under the recessive genetic model with homozygote relative risk *f*_2_/ *f*_0_ = 2.25 and with a risk allele frequency of *p* = 0.19, NCP_RA joint_ = 46.49 and NCP_genotypic_ = 42.76, while the NCP to achieve 80% power at *α* = 5 × 10^−8^ is 43.02. Figures 4e and 4f also show that, under the heterozygous advantage and heterozygous disadvantage genetic models, the difference between NCP_RA joint_ and NCP_genotypic_ can be substantial as the effect size increases.

**Figure 4:**
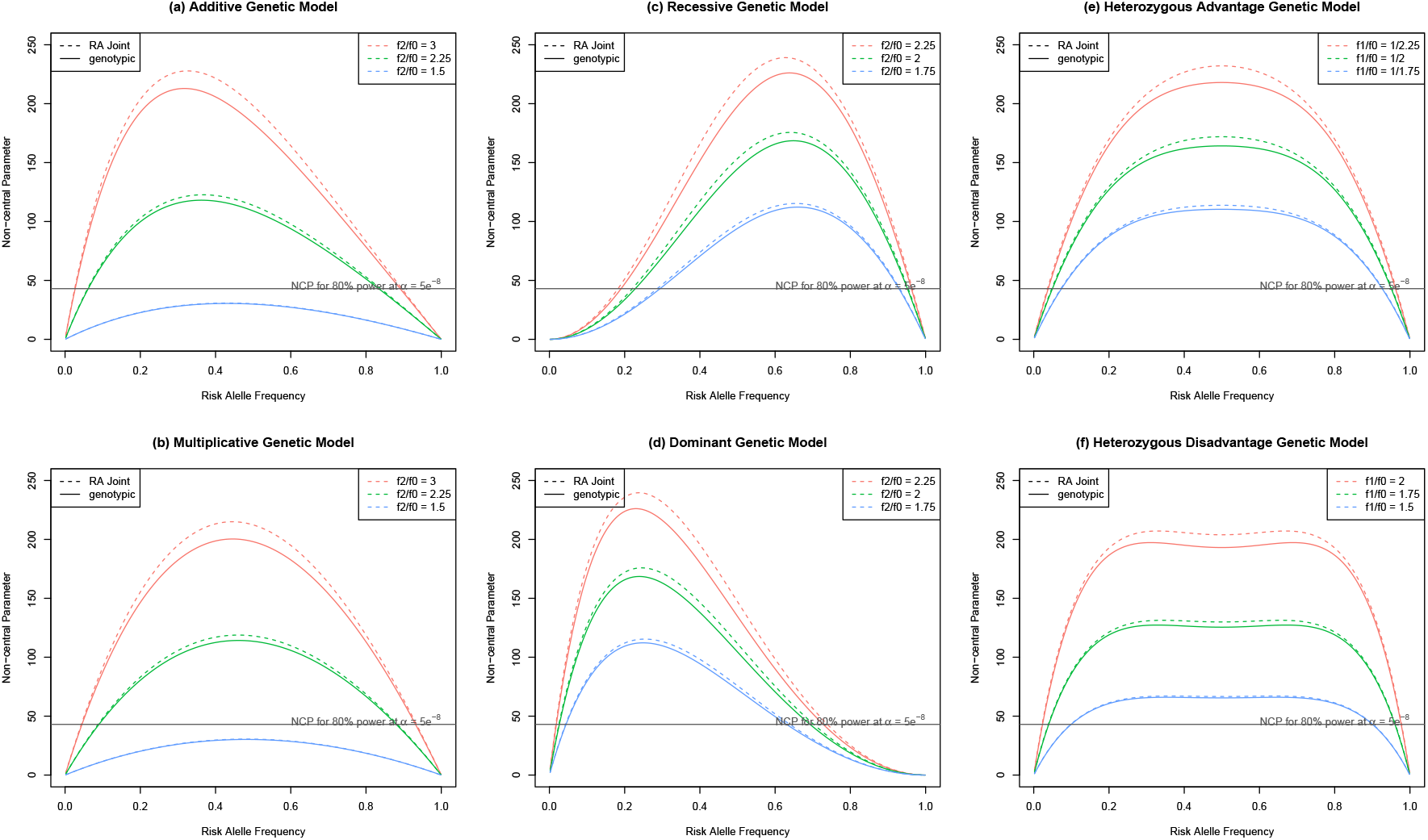
NCP_RA joint_ and NCP_genotypic_ under six common genetic models. The NCPs are computed assuming *r* = *s* =2,500, and HWE holds at the population level. *f*_2_ is fixed at 0.1. For the additive and multiplicative genetic models, the homozygote relative risk (*f*_2_/ *f*_0_) takes values from {3, 2.25, 1.5}; for the recessive and dominant genetic models, the homozygote relative risk ( *f*_2_/ *f*_0_) takes values from {2.25, 2, 1.75}; for the heterozygous advantage genetic model, the heterozygote relative risk ( *f*_1_/ *f*_0_) takes values from {1/2.25, 1/2, 1/1.75}; for the heterozygous disadvantage genetic model, the heterozygote relative risk ( *f*_1_/ *f*_0_) takes values from {2, 1.75, 1.5}. Dotted lines represent NCP_RA joint_ and solid lines represent NCP_genotypic_. The horizontal line represents the NCP magnitude necessary to achieve 80% power at *α* = 5 × 10^−8^ for a 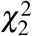 test.

## 8 Application Studies

### 8.1 Application 1 - the case-control studies of Wittke-Thompson et al. (2005)

We further demonstrate the merits of the proposed RA joint test by revisiting the 60 SNPs reported in Wittke-Thompson et al. (2005); these SNPs were studied by the authors to verify if HWD in the case or control sample is consistent with the underlying genetic models. Our analysis focuses on the 40 SNPs with all genotype counts ≥ 5 as tests can be sensitive to SNPs with low genotype counts. We adopt a conservative Bonferroni correction at *α* = 0.05/40 = 0.00125 to adjust for multiple hypothesis testing.

Figure 5 plots the −log_10_ p-values of (a) the traditional 1 d.f. additive test against the proposed RA joint test and (b) the traditional 2 d.f. genotypic test against the proposed RA joint test. For the three significant SNPs identified by all three tests, the p-values of the proposed RA joint tests can be one fold smaller than those of the other two tests. Note that this does not imply that the RA joint test always produces smaller p-values than the traditional additive test, since most of the SNPs studied here appear not to follow the additive genetic model (Wittke-Thompson et al., 2005). Compared with the traditional 2 d.f. genotypic test, the proposed RA joint produces smaller p-values for all the three significant SNPs, which validates our findings in Section 6.3 (simulated power comparison) and Section 7 (theoretical power comparison).

**Figure 5:**
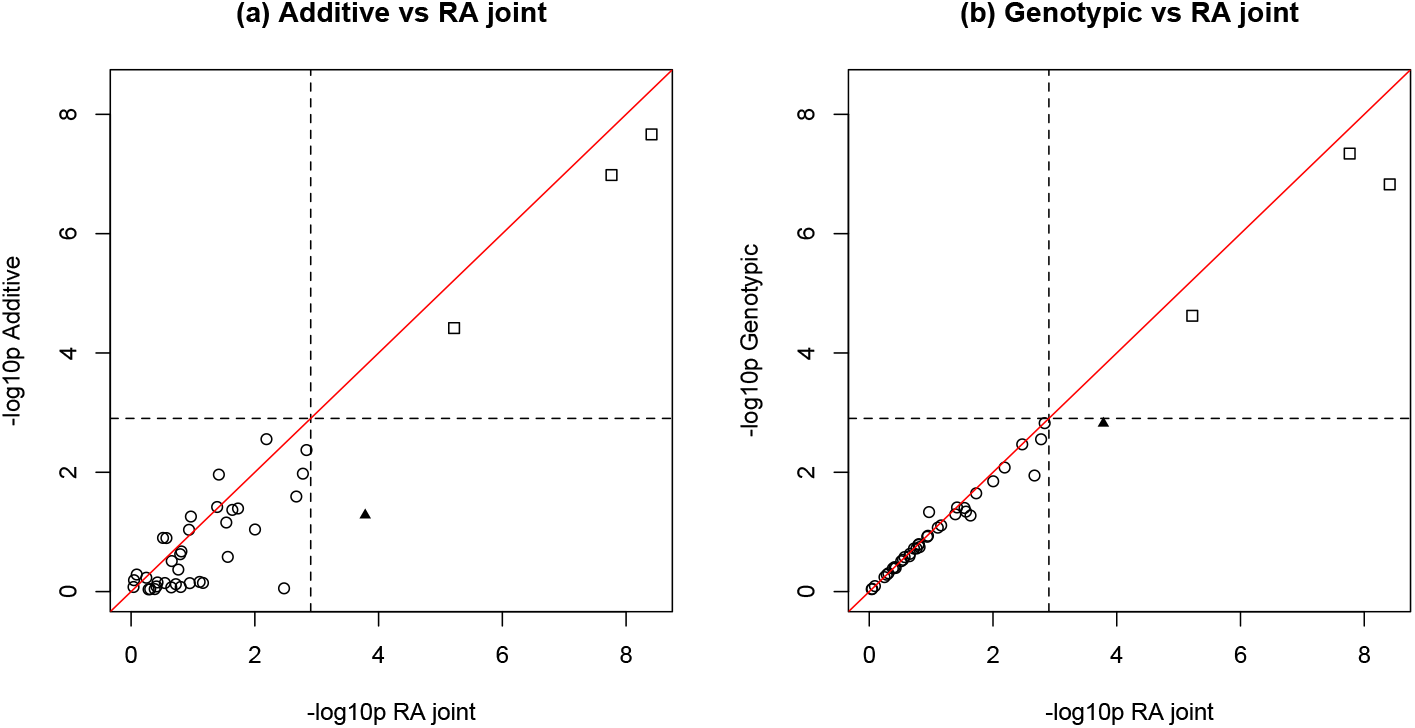
Results of application 1 - the case-control studies of Wittke-Thompson et al. (2005), comparing (a) the traditional additive test vs. the proposed RA joint test, and (b) the traditional genotypic test vs. the proposed RA joint test. The nominal value is *α* = 0.00125 correcting for multiple hypothesis to control the family-wise error rate (FWER) at 0.05. The squares mark the association tests that are significant using all three tests, and the solid triangle marks the association test that is only significant using the proposed RA joint test.

Noticeably, the association test between the ACE genotype and nephroangiosclerosis (marked by the solid triangle) is only significant using the proposed RA joint test at *α* = 0.00125. This is because the difference in HWD between the case and control samples 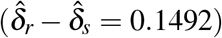 is sub-stantial compared to the difference in risk allele frequency 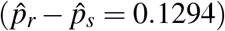. But the difference in HWD is only captured by the proposed RA joint test; see Table 6 for details.

**Table 6:** A case-control study of the ACE genotype and nephroangiosclerosis analyzed in application 1, demonstrating that the RA joint test utilizes the difference in HWD between the case and control groups to increase the power of detecting the association.

| | *aa* | *Aa* | *AA* | Total | $\hat{p}$ | $\hat{\delta}$ | $p_{\text{HWE}}$ | $p_{\text{RA joint}}$ | $p_{\text{genotypic}}$ | $p_{\text{additive}}$ |
| --- | --- | --- | --- | --- | --- | --- | --- | --- | --- | --- |
| Case | 6 | 10 | 21 | 37 | 0.7027 | 0.0737 | 0.0317 | — | — | — |
| Control | 8 | 48 | 19 | 75 | 0.5733 | −0.0754 | 0.0076 | — | — | — |
| Total | 14 | 58 | 40 | 112 | 0.6161 | −0.0224 | 0.3162 | $1.64 \times 10^{-4}$ | 0.0015 | 0.0522 |

## 9 Discussion

In this work, we introduce the RA joint test that simultaneously captures the difference in risk allele frequency and the difference in HWD measure between the case and control groups, yet robust to HWD not attributed to true association. We show that the proposed RA joint test is robust to HWD at the population level while incorporating the difference in HWD between the case and control samples to increase the power of case-control association test.

To examine the power of the proposed RA joint test, we conduct power studies under six common genetic models, and compare with the traditional additive association test and the traditional genotypic association test. Under the additive genetic model, the RA joint test, as expected, loses power over the traditional additive test. However, as in another testing context, Chen et al. (2018) has shown that the power difference between a 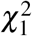 and a 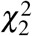 test statistic can be capped at 11%. We also show that this loss decreases as the population level HWD, *δ*, increases; see Supplementary Figure **??** for an example when *δ* = 0.04. Under the additive on log-scale genetic model, or the multiplicative genetic model, the power exhibits a similar pattern to that of the additive genetic model when the risk homozygote penetrance probability and the homozygote relative risk are the same in the two models. Under the recessive and dominant models, the proposed RA joint test gains slightly higher power over the traditional additive test, while under the heterozygous advantage and disadvantage models, the RA joint test gains substantial power over the traditional additive test. The latter happens when the risk allele frequencies between the case and controls are similar, but the genotype distributions hence HWD estimates are drastically different. This scenario is not only theoretically possible but also practically observable as demonstrated in Tables 6 and **??**.

In the empirical power studies, it appears that the proposed RA joint test is practically identical to the traditional genotypic test. We thus further compare the theoretical power between the two tests (both follow the 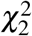 distribution) via non-central parameters. We show that the non-central parameter of the proposed RA joint test is larger than that of the traditional genotypic test under all six genetic models in balanced case-control studies. When the effect size is small, the difference between the NCPs of the two tests are marginal, but NCP_RA joint_ is slightly higher than NCP_genotypic_; see Supplementary Figure **??**. As the effect size increases, the difference in NCPs between the two tests can be substantial, indicating that *T*_RA joint_ can be more powerful to detect truly associated SNPs in GWAS, which is also verified by the application studies.

In the simulated and theoretical power studies, we have assumed equal numbers of cases and controls, i.e. *π*_01_ = *s/r* = 1. However, in real-life genetic association studies, often there are more controls than cases (i.e. *π*_01_ > 1). We show that the effect of *π*_01_ on NCP_RA joint_ and NCP_genotypic_ depends on the genetic models, risk allele frequencies and the disease penetrance probabilities; see Supplementary Figure **??** for details.

When the frequency of the risk allele is close to 0 or 1, e.g. *p*_*A*_ < 0.05 or *p*_*A*_ > 0.95, the corresponding SNP (often called rare variant) is likely to be removed from the analysis as the RA joint test tends to be anti-conservative for SNPs with low genotype counts. A study of rare variants is beyond the scope of our work, and we refer readers to Derkach et al. (2014) for a review of statistical methods developed for jointly analyzing multiple rare variants. However, here we make two remarks about the two 2 d.f. tests on analyzing one single rare variant. Considering a rare disease where the risk allele is the minor allele, in theory, the disease penetrance probability for the common homozygous genotype can be zero, i.e. *f*_0_ = 0, which implies no individuals in the case group have genotype *aa*, that is, *r*_0_ = 0. In this case, the traditional genotypic Wald’s test statistic goes to infinity without small-sample adjustment. Although the current implementation of the RA joint test filters out low-count SNPs, the test itself is capable of producing meaningful test statistics.

By convention, allele *A* is usually referred as the minor allele (i.e. *p*_*A*_ ≤ 0.5) with the assumption that diseases are rare, and it is the minor allele that increases the risk. In this work, we allow *p*_*A*_ to be bigger than 0.5, and broaden the notion of ‘disease’ traits to any rare or common phenotype traits. For example, in the current genetic studies of susceptibility and disease severity of COVID-19, the risk allele can be a major allele for disease susceptibility which is common; the risk allele is likely to be the minor allele for disease severity because it is rare (Lin et al., 2020). Therefore, in the simulated and theoretical power studies of this work, the range of *p*_*A*_ is from 0.05 to 0.95.

From this whole range of *p*, we observe the power of the proposed RA joint test, the traditional 1 d.f. additive test and 2 d.f. genotypic tests is not symmetric to *p* = 0.5 under additive, multiplicative, recessive, and dominant genetic models. Furthermore, the power of the tests are not always maximized at *p* = 0.5 as commonly believed‥ The power of all the tests is a complex function of the risk allele frequency whose shape and peak are determined by the penetrance probabilities.

In this work, we have laid the ground work for jointly testing allele frequency difference and HWD difference between the case and control samples to improve the power of the traditional association tests. To provide closed-form solutions for method comparison, we did not include covariates (e.g. age and sex) in our studies, but the proposed method can easily account for covariate effects. The proposed *allele-based* regression framework also opens up the possibility of analyzing more complex data such as pedigree data of related individuals, which we will explore as future work.

## References

Anderson, C. A., F. H. Pettersson, G. M. Clarke, L. R. Cardon, A. P. Morris, and K. T. Zondervan, 2010: Data quality control in genetic case-control association studies. Nature protocols, 5 (9), 1564–1573.

Apple, R. J., H. A. Erlich, W. Klitz, M. M. Manos, T. M. Becker, and C. M. Wheeler, 1994: Hla dr– dq associations with cervical carcinoma show papillomavirus–type specificity. Nature genetics, 6 (2), 157–162.

Bycroft, C., and Coauthors, 2018: The uk biobank resource with deep phenotyping and genomic data. Nature, 562 (7726), 203–209.

Chen, B., R. V. Craiu, L. J. Strug, and L. Sun, 2018: The x factor: A robust and powerful approach to x-chromosome-inclusive whole-genome association studies. arXiv preprint arXiv:1811.00964.

Derkach, A., J. F. Lawless, L. Sun, and Coauthors, 2014: Pooled association tests for rare genetic variants: a review and some new results. Statistical Science, 29 (2), 302–321.

Dudbridge, F., and A. Gusnanto, 2008: Estimation of significance thresholds for genomewide association scans. Genetic Epidemiology: The Official Publication of the International Genetic Epidemiology Society, 32 (3), 227–234.

Lin, Y.-C., and Coauthors, 2020: On statistical power for case-control host genomic studies of covid-19. medRxiv.

Marees, A. T., H. de Kluiver, S. Stringer, F. Vorspan, E. Curis, C. Marie-Claire, and E. M. Derks, 2018: A tutorial on conducting genome-wide association studies: Quality control and statistical analysis. International journal of methods in psychiatric research, 27 (2), e1608.

Sasieni, P. D., 1997: From genotypes to genes: doubling the sample size. Biometrics, 1253–1261.

Schaid, D. J., and S. J. Jacobsen, 1999: Biased tests of association: comparisons of allele frequencies when departing from hardy-weinberg proportions. American Journal of Epidemiology, 149 (8), 706–711.

Song, K., and R. C. Elston, 2006: A powerful method of combining measures of association and hardy–weinberg disequilibrium for fine-mapping in case-control studies. Statistics in Medicine, 25 (1), 105–126.

Taylor, J., and R. Tibshirani, 2006: A tail strength measure for assessing the overall univariate significance in a dataset. Biostatistics, 7 (2), 167–181.

Wang, J., and S. Shete, 2008: A test for genetic association that incorporates information about deviation from hardy-weinberg proportions in cases. The American Journal of Human Genetics, 83 (1), 53–63.

Wittke-Thompson, J. K., A. Pluzhnikov, and N. J. Cox, 2005: Rational inferences about departures from hardy-weinberg equilibrium. The American Journal of Human Genetics, 76 (6), 967–986.

